# Retention of a female-specific growth hormone receptor gene correlates with reverse sexual size dimorphism in birds

**DOI:** 10.64898/2026.04.10.717760

**Authors:** Shauna A. Rasband, Michael J. Braun, Philip L. F. Johnson

**Author notes:** **Author Emails:** Shauna A. Rasband Michael J. Braun Philip L. F. Johnson.

## Abstract

In bird species, larger males are the most common, followed by equal-sized sexes; larger females are rarest. Although multiple forces have been proposed to select for larger female body size, the molecular mechanisms determining this trait remain uncharacterized. The avian gene for growth hormone receptor (*GHR*) on the Z chromosome is a known factor in sex-specific development. Male birds (ZZ) have two copies of this gene, while females (ZW) typically have one. In addition to the Z copy of *GHR* found in all birds, we identified a second copy of *GHR,* documenting its presence or absence across the avian phylogeny by searching over a thousand genomes. This copy was solely found in females. When this second copy can be assigned to a chromosome, it is always located on the W. We infer that this second copy is a gametolog that was present in the common ancestor of extant birds, but has been lost from most lineages. *GHR-W* has notably been retained in hawks, eagles, vultures (Accipitriformes) and owls (Strigiformes). These groups display larger female size, making *GHR-W* an intriguing potential contributor to reverse sexual dimorphism. In these lineages, *GHR-W* retains intact gene structure, despite W-chromosome degradation that leads to loss of this gene from most birds. Analysis of published RNA-seq data demonstrates that *GHR-W* is expressed in accipitriform fibroblasts. Using available sex-specific mass data and correcting for phylogenetic structure, we report a robust correlation between retention of *GHR-W* and reverse sexual size dimorphism across avian clades.

## 1 INTRODUCTION

Sexual size dimorphism is a pervasive phenomenon in vertebrate taxa, with some lineages favoring larger males (e.g. mammals, birds) and others favoring larger females (e.g. amphibians, snakes and lizards) (Slavenko et al., 2025). While a majority of avian taxa have larger-bodied males, some lineages, including many birds of prey, have larger-bodied females (Székely et al., 2007). The disparity in body size is extreme in some cases; for example, female Harpy Eagles (*Harpia harpyja*) weigh 1.8x more than males (Dunning, 2007). Due to its rarity and sporadic appearance across the avian phylogenetic tree, the causes and implications of this “reverse” sexual dimorphism (RSD) have been longstanding matters of interest to the ornithological, evolutionary, and ecological science communities. The extensive early literature on this subject was summarized by Andersson (1994, pp. 275-284).

Both natural and sexual selection are thought to drive sexual size dimorphism. Natural selection on foraging success is one mechanism hypothesized to drive disruptive selection on body size. For example, intraspecific competition is reduced when the larger sex hunts larger prey and the smaller sex hunts smaller prey (Bauld et al., 2022; Zavalaga et al., 2007). This could result in RSD, though it could equally select for male-biased sexual size dimorphism (SSD) or even intraspecies size polymorphism unrelated to sex. However, sex-specific resource-gathering and reproductive roles may predispose a clade towards RSD, as in raptors where nesting females brood young and males hunt prey. If smaller body size increases hunting efficiency due to higher agility, then small males would be selected for, whereas female body size would not be under this selective pressure (Slagsvold & A. Sonerud, 2007; Sonerud et al., 2014). Additionally, larger female body size lessens the magnitude of weight fluctuation during the gravid period and thus may be under positive selection to maintain female foraging success while carrying unlaid eggs.

Foraging success is more variable in species that pursue large, scattered prey. In fact, hunting this type of challenging prey is highly correlated with both the presence and magnitude of RSD in birds of prey (Krüger, 2005). Natural selection on fecundity also favors larger females, which can make greater energetic investments in more eggs. This is essentially Darwin’s fecundity advantage hypothesis (Darwin, 1871), which has been extensively borne out in the literature in poikilothermic organisms (Preziosi et al., 1996; Blanckenhorn, 2005). For homeothermic animals like birds, a variant of Darwin’s fecundity advantage hypothesis is sometimes postulated with higher energetic investment via higher-quality offspring, rather than an increased number (Blanckenhorn, 2005). However, the evidence that fecundity advantage drives RSD in birds is inconsistent (Pincheira-Donoso & Hunt, 2017). Lastly, in certain taxa, sexual selection may play a role, when females choose small males who perform the best acrobatic flight displays (Blomqvist et al., 1997; Shogren et al., 2022). Despite the multifarious previously hypothesized selective forces that may drive this growth pattern, the genetic underpinnings of RSD remain unexplored.

In birds, body size is regulated by the pituitary-hypothalamic axis, including the growth hormone receptor (GHR), which is encoded by a sex-linked gene (*GHR*) 132 kb in length on the Z chromosome. In contrast to mammals, which typically have a XY sex system, birds have a ZW sex system; male birds are homogametic, possessing two Z chromosomes, whereas female birds are heterogametic, with one Z and one W chromosome. As neither copy of the Z chromosome is uniformly silenced in male (ZZ) birds (Wolf & Bryk, 2011), the gene dosage of many Z-linked loci for males is twice that of females (ZW). Thus, the Z chromosome location of *GHR* may result in comparatively lower expression of this gene in many female (ZW) birds, as observed in a recent study in a model passerine (Davenport et al., 2023). Sufficient expression of *GHR* is needed for normal growth processes and cell proliferation; e.g., body growth is highly correlated to *GHR* expression in liver tissue in chickens (H. Y. Huang et al., 2016). In response to growth hormone binding, GHR in the liver causes the secretion of insulin-like growth factor 1 (IGF-1). In vertebrates, circulating IGF-1 drives whole-body growth, mainly via proliferation of cells in cartilage, muscle, and bone (Brooks & Waters, 2010; Dantzer & Swanson, 2012). It is therefore possible that the sex-linked *GHR* could be involved in sexual size dimorphism in birds.

We previously reported the existence of a duplicate growth hormone gene (*GH*) in passerine birds (Yuri et al., 2008) and characterized its presence across all major passerine lineages (Rasband et al., 2023). The passerine *GH* duplication inspired us to search for duplications of the receptor gene, *GHR*, as both synchronous and asynchronous duplications of the *GH-GHR* receptor-ligand gene pair have occurred in other vertebrate lineages (Ocampo Daza & Larhammar, 2018). While most bird genomes have a single copy of the *GHR* gene on the Z chromosome, our searches of genomic data beyond passerines revealed that a small subset of non-passerine birds have a second copy of *GHR,* found exclusively in female genomes. When duplicate copies could confidently be assigned to a chromosome, they were found on the female-limited W sex chromosome. We refer to these W-linked copies as *GHR-W*.

Here we characterize the phylogenetic distribution of duplicate *GHR* genes in birds and establish the genomic origins of the *GHR-W* gene. We show that both *GHR-Z* and *GHR-W* are expressed in relevant transcriptome data and that the presence of duplicate *GHR* genes is strongly correlated with RSD across the avian tree of life.

## 2 MATERIALS AND METHODS

### 2.1 *GHR* Sequence Curation

We used blastn (Altschul et al., 1990) to search for *GHR* genes in avian genomes available in GenBank/RefSeq as of February 29, 2024. We analyzed a maximum of one genome per species per sex for a total of 1104 avian genomes from 1078 species. In our blast searches, we used the full length chicken (*Gallus gallus*) *GHR-Z* mRNA (NCBI: NM_001001293.2) as our query sequence. Although chicken is phylogenetically distant from many bird species, we expected that we would still find *GHR* sequence hits in many taxa as this gene is well-conserved across vertebrates. Indeed, we found *GHR* sequence hits for most genomes, including those of birds belonging to the most distantly related groups (Neoaves and Paleognathae).

We assessed the strength of evidence of multiple *GHR* copies in each genome. At times, genome assembly can result in artifactual duplication where one gene copy is incorrectly assembled on multiple scaffolds. To avoid these false positives, we defined evidence of multiple *GHR* copies as multiple sequence hits that 1) overlapped on the query sequence and 2) had multiple sequence differences in the overlapping region. If genomes did not meet these two criteria, we defined them as having “no evidence” of multiple copies of *GHR.* We categorized evidence of multiple copies of *GHR* as “good” if two hit sequences each had >50% query coverage and “limited” if one or both hit sequences had <50% query coverage (Supplemental Table 1). Genomes that returned no hits in our blast searches were too incomplete for further analyses (also listed in Supplemental Table 1). To explore the phylogenetic distribution of multiple copies of *GHR* across birds, we mapped the proportion of genomes showing “good,” “limited,” and “no evidence” of multiple *GHR* copies onto a pre-existing phylogenetic tree (Braun et al., 2024), which includes all deep avian lineages.

### 2.2 Determining Genomic Sex

We collected all available genomic sex metadata as listed directly on NCBI or attached to museum accession codes for the voucher specimens sequenced. We also inferred female sex when an assembled W chromosome was present, and from personal communication with the genome submitter (pers. comm., Catanach T., 2023). However, many genomes lacked sex data from these sources. We determined the sex of certain genomes where the sex annotation was suspect or not reported, listed in Supplemental Table 2—along with the reason for determining the genomic sex in each case. As there were hundreds of genomes lacking sex data, we focused on a subset to accomplish the following aims: 1. To check if all genomes with any evidence of two copies of *GHR* are female; 2. To assess the extent of two copies of *GHR* in Accipitriformes by A) determining the sex of accipitriform genomes with unknown sex, and B) by checking the sex of accipitriform genomes listed as female but which lacked evidence of 2 *GHR* copies. We also sexed five known male genomes in relevant clades to assess the spread of Z:A coverage ratios in this sex.

To calculate the Z:A coverage ratio to determine genomic sex, we obtained the raw reads for such genomes from the NCBI Sequence Read Archive and mapped them to a related chromosome-level reference genome using the mem tool of bwa version 0.7.17 (Li & Durbin, 2009) with default settings. We used samtools version 1.17 (Danecek et al., 2021) to process the bwa output: converting the sam output file to bam (samtools view), sorting and indexing that file (samtools sort, samtools index) and then obtaining the read depth (samtools depth) of the Z chromosome and chromosome 1, an autosome of similar size. In theory, males should have a Z to autosome (Z:A) coverage ratio of ∼1, whereas females should have a Z:A coverage ratio of ∼0.5.

In practice, the mean Z:A ratio across samples showed more variation than we expected (Supplemental Figure 1; Supplemental Table 2), which we hypothesized was due to some W reads incorrectly mapping to Z as a result of the incomplete nature of many assembled sex chromosomes in the reference genomes. Indeed, when we analyzed the read coverage in non-overlapping windows of 100 kb over the Z chromosome, for female genomes we saw two distinct coverage ratios (∼1 and ∼0.5) when dividing the Z coverage level by the mean coverage level of the comparison autosome. These coverage levels occurred in different regions of the Z chromosome (Supplemental Figure 2A), whereas confirmed male genomes had just one ratio (∼1) that did not vary across the Z (Supplemental Figure 2B). We therefore designated those genomes with two distinct levels of Z and W chromosome coverage as female (mean ratio 0.52 to 0.86) and those with one level as male (mean ratio 0.95 to 1.27) as reported in Supplemental Table 2.

### 2.3 Sequence Alignment and Gene Tree Estimation

To estimate an avian *GHR* gene tree, we subsampled representative genomes with evidence of multiple *GHR*s from across the avian phylogeny. Since many genomes have incomplete *GHR* loci due to low-coverage Z/W scaffolds, we used sequence data from the portion of *GHR* that was most frequently intact: a 4-6 kb region with the highest exon density in the gene, spanning the final three exons as well as the 3’ untranslated region, or UTR. This portion corresponds to exons 8-10 and introns 8-9 of chicken *GHR* transcript variant 1 (NM_001001293.2). Lastly, we included the chicken *GHR-Z* sequence as a representative from a well-assembled Neognath genome without any evidence of multiple *GHR* copies. Because pseudogenization is difficult to assess in partial or gapped sequences, we did not assess *GHR* gene integrity for the purpose of tree building. To multiply align the selected sequences, we used MAFFT v7.490 (Katoh & Standley, 2013) with standard settings (Algorithm: Auto, selecting the best strategy for the underlying sequences, scoring matrix 200PAM / k=2, gap open penalty of 1.53 and offset value of 0.123). We then estimated a maximum likelihood gene tree using IQ-TREE v2.3 (Nguyen et al., 2015) on standard settings (substitution model: auto, search parameters: 0.5 perturbation strength and IQ-TREE stopping rule at 100) and assessed nodal support with 1000 bootstraps.

To evaluate whether the region surrounding *GHR-Z* and *GHR-W* genes may lie in the pseudo-autosomal region (PAR) and may still be recombining, we examined sequence similarity in ∼1 megabase regions flanking *GHR-W* and *GHR-Z* of high-quality exemplar genomes. We chose Eurasian Goshawk (*Astur gentilis,* GCA_929443795.1) and Elegant Crested-Tinamou (*Eudromia elegans*, GCA_047922985.1) genomes to represent Neoaves and Paleognathae, respectively. Both are chromosome-level genome assemblies with both *GHR-Z* and *GHR-W* full length and apparently intact (maintaining exon/intron gene structure and open reading frame).

Centering on the *GHR-Z* and *GHR-W* genes, we excerpted 500 kb of DNA upstream of the first exon and 500 kb downstream of the last exon of each. We compared these regions in a dotplot with dgenies using Minimap2 v2.26 (Li, 2018) to visually evaluate sequence similarity in these regions (Figure 3).

**Figure 1:**
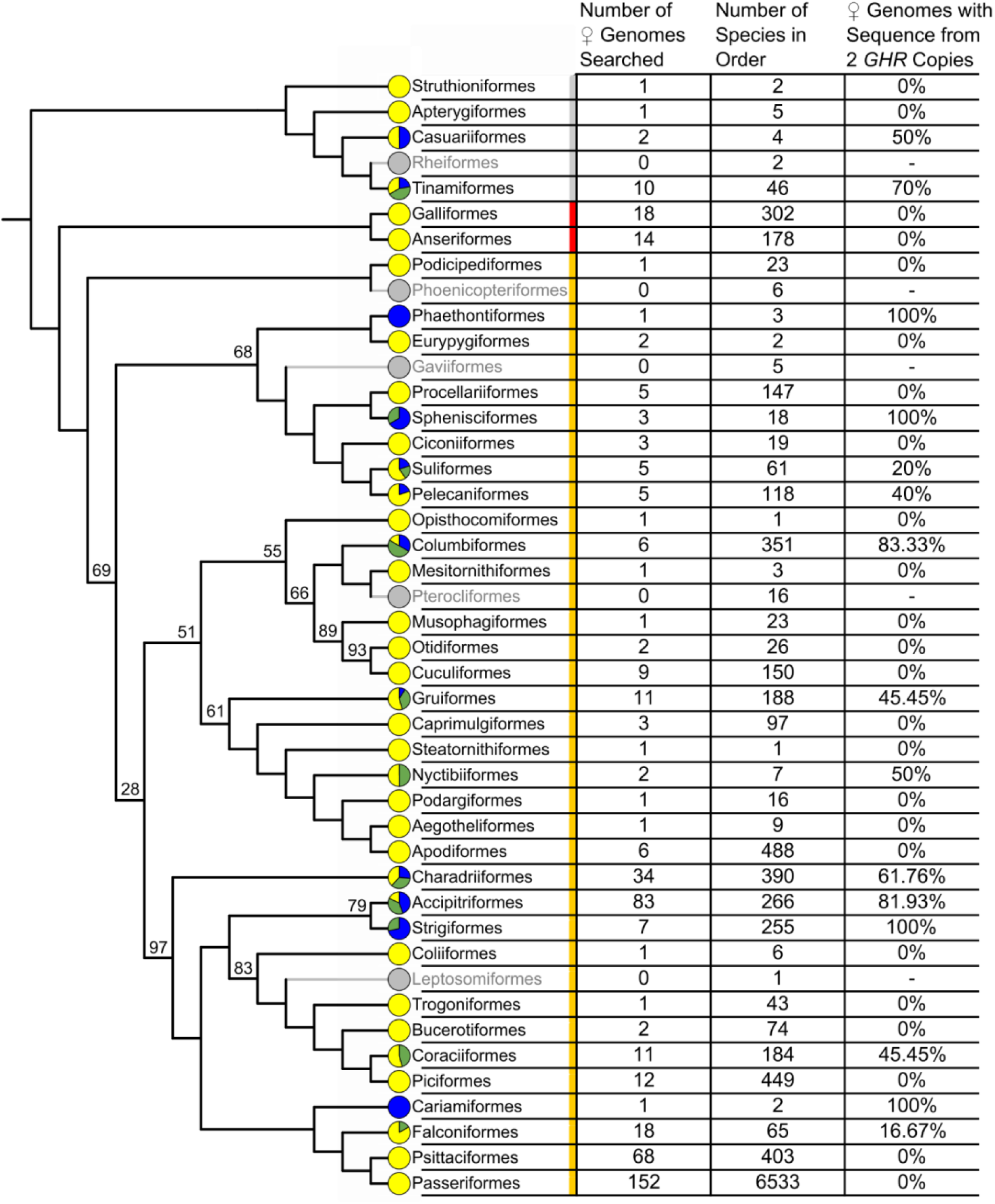
An avian ordinal phylogeny (derived from Braun et al., 2024, figure 2) onto which is mapped the presence/absence of two copies of GHR in genomes derived from female individuals. Where bootstrap values are less than 100, they are shown on the relevant nodes. Pie charts on each tip represent the prevalence of evidence of two copies of GHR in female genomes sampled from that order. On the pie chart, species with good evidence of two copies are marked in dark blue (the darkest color), species with limited evidence of two copies are marked in green, and species with no evidence of two copies are marked in yellow (the lightest color). Missing data, i.e. avian orders without female genomes, are marked with paler gray branches leading to gray pie charts. Ordinal names are shown for each branch and supraordinal clades are shaded: gray for Paleognathae, red for Galloanserae, and goldenrod for Neoaves. The table to the right shows the number of female genomes searched, the number of species in each order according to Gill et al. (2024), and the percentage of female genomes searched that had any sequence from two copies of GHR, i.e. genomes with good evidence of two copies of GHR plus genomes with limited evidence of two copies of GHR.

**Figure 2:**
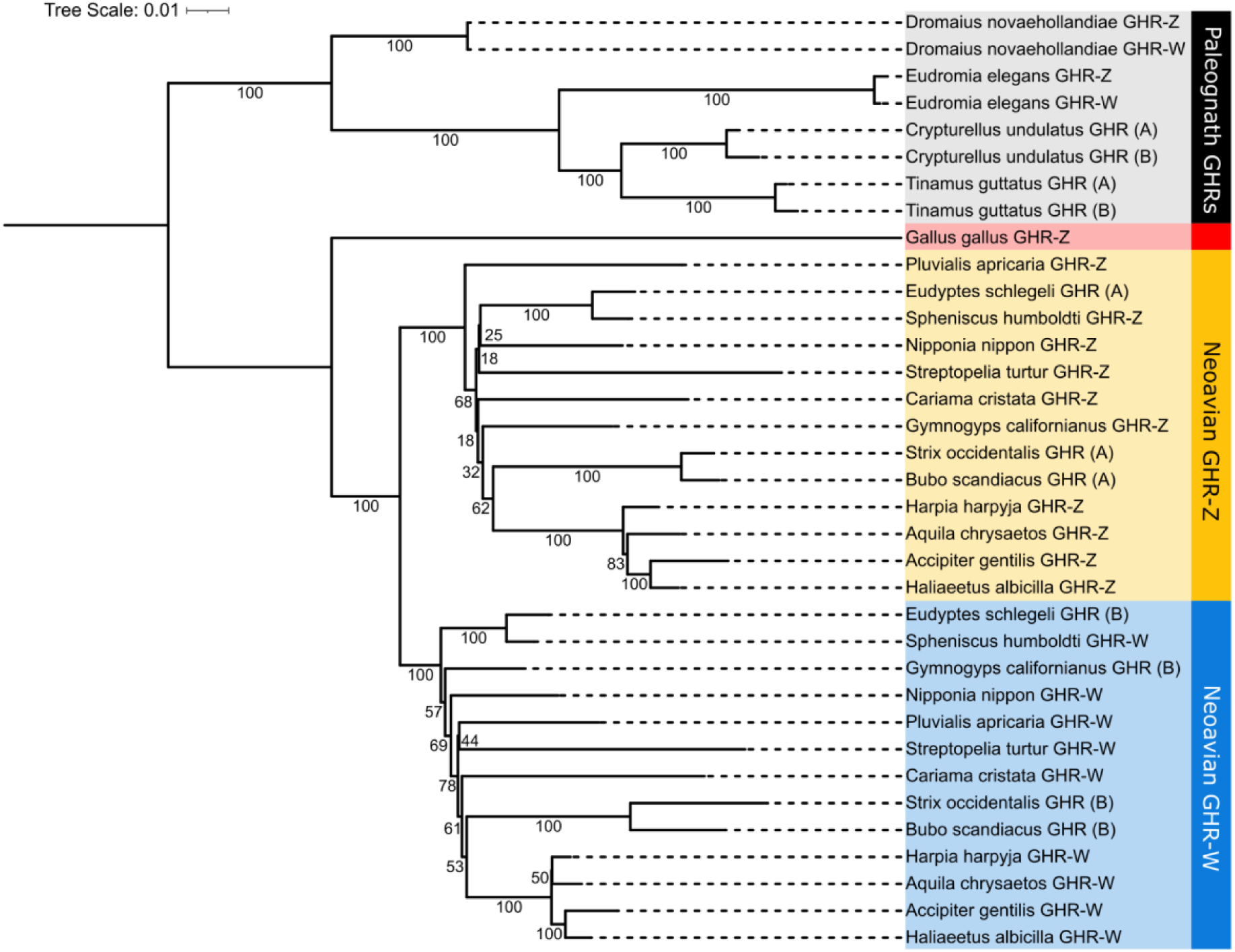
Maximum likelihood tree of avian GHR genes (see Methods 2.3 for details). The tree is rooted with Paleognaths as the known outgroup. Leaves are labeled GHR-Z or GHR-W if found on assembled sex chromosomes. GHRs from genomes without full chromosome-level assemblies are labeled A or B. Major clades are marked with colored vertical bars on the right: black, Paleognathae; red, Galloanserae GHR-Z; goldenrod, Neoaves GHR-Z; blue, Neoaves GHR-W. Branch labels show nodal support values, calculated as a percentage of 1000 bootstraps.

**Figure 3:**
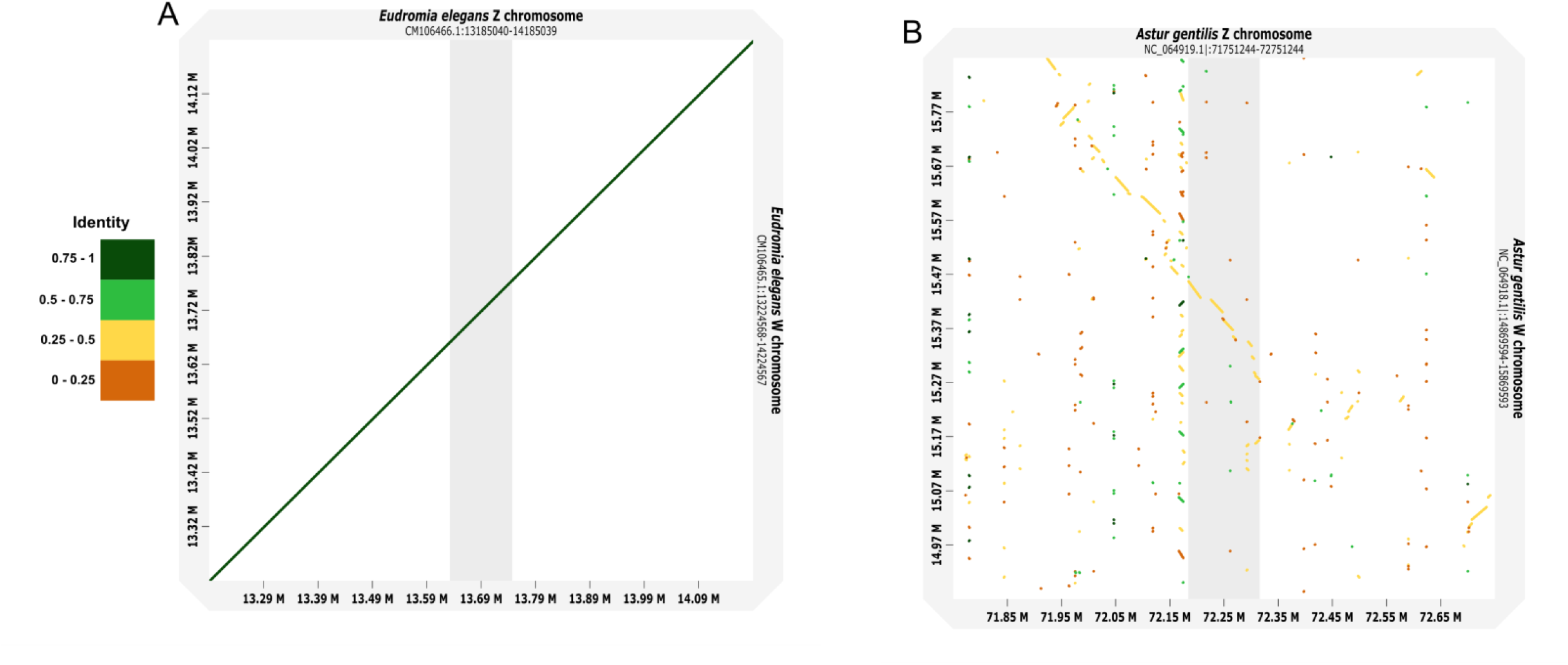
Dotplots centered on GHR-Z and GHR-W showing 500 kb on either side of the gene, aligned with dgenies using Minimap2 v2.26. Sequence identity is color coded and calculated as M/N, where M is the number of matching bases in the mapping and N is the total number of bases, including gaps. The portion corresponding to GHR-Z is highlighted in gray. **3A:** Eudromia elegans (genome: GCA_047922985.1). **3B:** Astur gentilis (genome: GCA_929443795.1).

### 2.4 Evaluating Evidence of *GHR-Z* and *GHR-W* Expression

To investigate *GHR* gene expression, we needed a species with a high-quality genome with two copies of *GHR* as well as female RNAseq data. We restricted our search to genomes with two copies of *GHR*, each of which had full-length open reading frames and canonical exon-intron structure with intact splice sites. The only species meeting these criteria as of February 29, 2024 was the California Condor (*Gymnogyps californianus*). Using STAR version 2.7.11b (Dobin et al., 2013), we aligned RNAseq reads from pooled fibroblast cells from two females (SRA: SRR17155298) to the California Condor genome assembled from the DNA of one these two females (GCA_018139145.2). As our use of default STAR settings allowed for multiple mismatches and multiple read alignment, we visualized and evaluated the specificity and uniqueness of reads mapped to each *GHR* gene in Geneious version 2022.2.1 (*Geneious Prime®*, 2022). We considered reads that mapped only to one copy (mapping quality value of 255 from STAR) as *GHR-Z* or *GHR-W* specific.

### 2.5 Testing the Correlation of Two *GHR* Copies to RSD

For all female genomes from which we retrieved at least one *GHR* sequence, we recorded the RSD status for that each species using mass as the metric representing size, with sex-specific mass data obtained from species accounts in *Birds of the World* (Billerman et al., 2024)*, The CRC Handbook of Avian Masses* (Dunning, 2007), and museum specimen records available on VertNet.org (institution codes: CAS, CHAS, CM, CRCM, CUMV, DMNS, KU, LACM, MCZ, MLZ, MSB, MVZ, NHMUK, ROM, TCWC, UAM, UBCBBM, UF, UMMZ, UMYMV, WFVZ, YPM). While most species had sex-specific mass data from numerous individuals, others had sex-specific mass data for only a few adult specimens, with additional qualitative data available, such as statements in *Birds of the World* that males are always heavier than females.

We therefore treated RSD as binary in this analysis, coding species with females larger than males as “RSD” and species with equal-sized sexes or larger males as “no RSD” (Supplemental Table 3). As phylogenies obtained from a single gene (e.g. *GHR*) may not accurately reconstruct avian relationships, for the analysis and data visualization, we used the multilocus species-level tree depicted in figure 4 of Braun et al., 2024, pruning taxa for which we could not find both female genomic and sex-specific mass data. Where genera represented on that tree had multiple species available with both types of data, we used a random number generator to select a maximum of two species per genus. We performed a phylogenetic generalized least squares logistic regression that accounts for phylogenetic structure in R version 2024.04.1+748, (R Core Team, 2021) using the package phylolm, version 2.6.2 (Tung Ho & Ané, 2014) to test the correlation of RSD with evidence of multiple *GHR* copies after correcting for phylogenetic structure. Both RSD and evidence of two copies of *GHR* were treated as categorical variables, either present or absent in each taxon.

**Figure 4:**
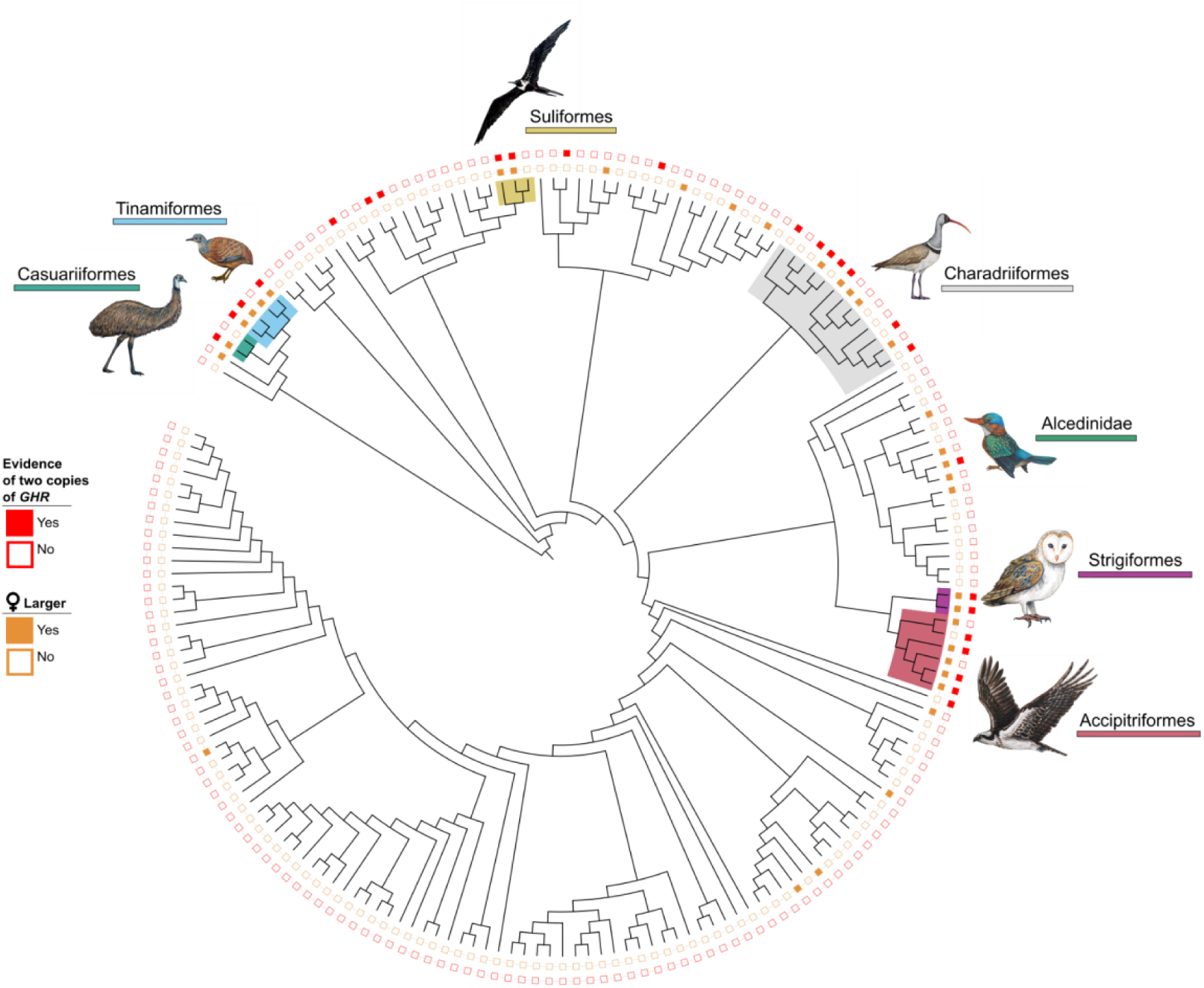
Mapped on a phylogeny of all extant bird groups derived from (Braun et al., 2024) are the character states of having two GHR genes and RSD. Outer ring: evidence of two GHR genes. Inner ring: the binary character state for RSD. Orders containing at least one species with both two copies of GHR and RSD are labeled and their clades shaded in the tree.

## 3 RESULTS

### 3.1 Avian Genomes with Evidence of Multiple *GHR* Copies

Of the 1104 avian genomes searched using BLAST, 1065 returned at least one hit for the chicken *GHR* query, and 133 of those yielded at least some evidence of two copies of *GHR* (Supplemental Table 1). Identifying a second copy of *GHR* proved difficult in some cases because these genomes spanned a wide range of assembly qualities. Moreover, the W chromosome is the most difficult chromosome to assemble because it is present only in females, is single-copy, and has high repeat content. Thus, the evidence for multiple copies of *GHR* in many genomes was partial or fragmentary.

For each genome, we categorized evidence for multiple copies as ‘good’ (64 genomes), ‘limited’ (69 genomes) or ‘no’ evidence (see Methods Section 2.1). In no case did we find evidence of more than two copies by our classification criteria. Of the 472 genomes previously identified as female or pooled female and male, 95 genomes had at least some evidence of two *GHR* copies (Supplemental Table 1). Contrastingly, of the 216 genomes recorded as male in the same data, only two had some evidence of two *GHR* copies (Supplemental Table 1).

To further investigate this apparent sexual asymmetry in *GHR* copy number, we performed genomic sex determination for the two “male” genomes with evidence for two copies as well as for the 36 genomes with evidence of two copies but no previous sex identification (see Methods: 2.2: Determining Genomic Sex). Our analyses revealed that the two “male” genomes with two *GHR* copies had sex mislabeled and were, in fact, derived from females (see Supplemental Figures 2C & 2D). Additionally, we found that 34 of the 36 unsexed genomes with evidence of two copies of *GHR* were female (2 remain unsexed as they lacked appropriate/available short reads for our method of sex determination). Therefore, none of the 229 male genomes we searched had any evidence of a second copy of *GHR*, while 131 of 506 female genomes searched had at least some evidence of two copies, and 2 unsexed genomes had some evidence of two copies. We did not attempt to determine the sex of the remaining unsexed genomes that lacked evidence of a second copy of *GHR,* except for twelve accipitriform genomes, to assess the extent of two copies of *GHR* in females in this clade.

### 3.2 Phylogenetic Distribution of Two *GHR* Copies

We used a previously published avian phylogeny (Braun et al., 2024; see methods) to broadly analyze the phylogenetic distribution of two *GHR* copies, which was uneven throughout the tree (Figure 1; Supplemental Table 1). We found evidence of two *GHR* copies in 13 of the 33 neoavian orders sampled (125 species total: 123 of 462 neoavian species examined, 460 with female genomes, and 2 with unsexed genomes). Paleognaths showed a distinctly higher proportion, with evidence of two copies in genomes from two of four orders (eight of fourteen genomes from thirteen paleognath species examined). Finally, we found no evidence of two *GHR* copies in either of the two galloanserian orders, even though these were relatively well sampled across 32 species with female genomes examined.

Of 133 species with evidence for two *GHR* copies, 68 were in the neoavian order Accipitriformes (hawks, eagles and New World vultures), the highest number of species in any order (Figure 1). Evidence of two *GHR* copies was also strong among owls (order Strigiformes), with two partial or full *GHR* sequences retrieved from all seven female genomes available. Of note, Accipitriformes and Strigiformes may well be sister groups, although the evidence for this relationship is not yet conclusive (see Stiller et al. 2024 for a recent analysis). Thus, this putative clade appears to represent an exception to the overall sporadic appearance of two *GHR* copies across neoavian orders. The fact that not all female accipitriform genomes showed evidence of two copies of *GHR* may be at least partly artifactual due to the draft nature of many genome assemblies; 100% of chromosome-level assembled genomes had good evidence of two copies of *GHR,* whereas only 41% of scaffold-level genomes did. Improved genome assemblies or targeted PCR assays could clarify whether any female accipitriforms are truly missing a second copy of *GHR.* A third order with a high percentage of taxa with two *GHR*s was Charadriiformes (shorebirds and relatives), with 21 of 34 female genomes displaying such evidence (Supplemental Table 1).

### 3.3 *GHR* Structure and Copy Location

Among avian *GHR* genes, that of chickens has been most thoroughly studied. It comprises 132 kb of the Z chromosome, with 10 exons, 9 introns (in transcript variant 1, NM_001001293.2) a long (194 bp) 5’UTR, and a 1,853 bp 3’ UTR. The coding sequence in this primary transcript is 1827 bp, encoding a polypeptide of 608 amino acids. In avian genomes we examined with two fully assembled *GHR* genes, this basic structure appears to be well conserved. Genomes of seven species spanning three orders (Accipitriformes, Casuariiformes, Strigiformes) had the gene structure of both *GHR* copies annotated in RefSeq (as *GHR* and *GHR-*like). These annotations showed copies on Z and W with conserved structure, including open reading frame, exon-intron structure, splice sites, and no premature stop codons (Supplemental Table 1).

Several lines of evidence indicate that in most or all avian genomes with two copies of *GHR,* the two copies are gametologs, i.e. located on the Z and W chromosomes. First, in all 13 chromosome-level genomes with two copies of *GHR,* one copy is always on Z and the other on W (with the possible exception of *Gymnogyps californianus* where the putative W copy is on a shorter scaffold that has not yet been assigned to a chromosome). These chromosome-level assemblies are broadly distributed across eight orders spanning the avian phylogeny (Accipitriformes, Cariamiformes, Casuariiformes, Charadriiformes, Columbiformes, Pelecaniformes, Sphenisciformes, Strigiformes), indicating that the Z and W genomic locations are widespread, if not universal, among birds with two copies of *GHR.* Second, in the 123 chromosome-level genomes (spanning 21 orders) with a single copy of *GHR*, *GHR* is located on the Z chromosome, further supporting a gametologous origin of the second copy present in some birds. Further, we found no evidence of tandem or Z to autosome duplication; all male genomes had one *GHR* copy only, and in all 120 scaffold-level assemblies with two copies of *GHR*, those two copies were assembled to separate scaffolds.

### 3.4 The Avian *GHR* Gene Tree

We sought to further elucidate the relationships between *GHR* sequences via tree building using both *GHR* sequences sampled across a breadth of avian orders (Figure 2; see Methods 2.3: Sequence Alignment and Gene Tree Estimation). In the resulting gene tree, there is a clear separation of neoavian *GHR* sequences into two clades: one containing all annotated Z chromosome copies and the other containing all annotated W chromosome copies. For those species without chromosome-level assemblies, one copy clusters strongly with each clade. These copies are labeled A in the clade with Z copies and B in the clade with W copies (Figure 2). The 100% bootstrap support for each of the two neoavian *GHR* clades owes to the substantial sequence divergence between them (mean 6.89% in CDS) and multiple fixed differences in both CDS and 3’ UTR (see alignment, Supplemental Data). In contrast, in paleognath species the two copies of *GHR* within a species always clustered together with minimal divergence between the two copies from the same taxon and substantial divergence among taxa.

One possible explanation for this difference between paleognath and neoavian *GHR* copy clustering is that, in paleognaths the copies may be allelic and recombining within species, while in neognaths they are not. Such a situation would arise if *GHR* copies in paleognaths are located in pseudo-autosomal regions (PARs) of sex chromosomes and neoavian *GHR*s lie outside of such regions. PARs are portions of the sex chromosomes that promote pairing during cell division and maintain sequence similarity due to recombination. To test this possibility, we examined sequence similarity in flanking regions near *GHR* genes of exemplar neognath and paleognath genomes. Dot plots revealed that sequence similarity was very high near the paleognath *GHR*s (Figure 3A), as expected if in a PAR, but much lower near the neognath *GHR*s, as expected if outside a PAR (Figure 3B). Patterns of *GHR* sequence conservation correspond to these example dotplots of regional sequence similarity; in paleognathous birds, nucleotide sequence conservation between Z and W copies of the coding sequence (CDS) is high (99.86% average sequence identity between fully assembled copies for *Dromaius novaehollandiae* and *Eudromia elegans*). In Neoaves however, the Z and W copies have diverged more in sequence (94.41% average sequence identity between annotated copies); for example, the Eurasian Goshawk (GCA_929443795.1) coding sequences have diverged by 6.21%, including 46 non-synonymous substitutions.

### 3.5 *GHR-W* Expression

As W copies of *GHR* may have been transcriptomically silenced or otherwise pseudogenized during the loss of W-chromosome content rampant in neoavian clades (Zhou et al., 2014), we aimed to investigate if both *GHR-Z* and *GHR-W* are transcribed in neoavian birds. Relevant transcriptome sequencing data in GenBank is limited, but we were able to map RNAseq reads from a California Condor (*Gymnogyps californianus*) fibroblast tissue culture to the chromosome-level genome assembly for the same species. Consistent with both copies being expressed, reads mapped across the full coding sequence of both gene copies. 47% of reads that mapped to the copy we infer is *GHR-W* based on the gene tree (Figure 2) were uniquely mapped (i.e. to divergent sequence areas) across this gene. Likewise, the reads mapping uniquely to the

*GHR-Z* copy spanned the gene, but were uniquely mapped at a much higher rate, 93%. The higher unique read-mapping rate and higher number of reads mapped to *GHR-Z* compared to *GHR-W* (15:1 ratio of unique *GHR-Z* reads to unique *GHR-W* reads) likely indicate that *GHR-Z* is expressed at higher levels than *GHR-W* in the source fibroblast tissue culture. However, given that cells from two individuals were pooled in the sample, we cannot infer individual expression levels of these two genes.

### 3.6 Correlation of *GHR-W* to RSD

In our phylogenetic trait mapping at the ordinal level (Figure 1), we found that many species with two copies of *GHR* are in clades noted for having larger females than males, such as Accipitriformes and Strigiformes. Motivated by the known role of GHR in body mass attainment in birds, we examined if this correlation of RSD with multiple *GHR* copies was confined to this clade or applied to birds in general. In figure 4, we visualize these variables on a species-level tree. The phylogenetic generalized least squares logistic regression model (accounting for phylogenetic structure) identified duplicate *GHR*s as strongly predictive of RSD (p < 0.000005). The pattern was driven by Charadriiformes, Accipitriformes, Casuariiformes, Suliformes, Strigiformes, Phaethontiformes, Tinamiformes, all of which had both RSD and two copies of *GHR* (Figure 4). Note that this correlation is not perfect, as Apterygiformes, Cuculiformes, and Falconiformes have RSD but no evidence for two *GHR*s, and Columbiformes, Gruiformes, Sphenisciformes, and Cariamiformes lack RSD but have some evidence for two *GHR*s.

## 4 DISCUSSION

The *GHR* gene in birds normally resides on the Z chromosome and thus exists at double the copy number in males as in females. We identified a second copy of *GHR* found in genomes of female birds distributed non-uniformly across the avian phylogeny (131 of 506 female genomes examined). Further investigation revealed that all chromosome-level assemblies showed the second copy to be on the W chromosome, bringing the copy number to parity between the sexes in these species. Gene tree analysis confirmed that, for the neoavian genomes with two copies sampled, one copy fell in a clade containing all annotated *GHR-W* genes while the other copy fell in a clade containing all annotated *GHR-Z* genes. Female paleognath genomes also exhibited copies on Z and W, but, in contrast to neoavians, the two copies were highly similar within each genome, consistent with allelic copies experiencing current or recent recombination during meiosis. These patterns suggest a scenario in which *GHR-W* originated not through duplication, but rather through retention on the W chromosome in some lineages, as avian sex chromosomes evolved from an ancestral autosome. The rampant reduction of gene content on the W chromosome then explains the absence of *GHR-W* from most avian genomes. Together, these observations raised the intriguing possibility that retention of *GHR-W* might be adaptive in some lineages.

### 4.1 Genomic Origin of *GHR* Genes in Birds

While *GHR* is autosomal in other vertebrates, it is sex-linked in birds. All extant birds share the same ZW sex chromosome pair, which was recruited prior to their diversification (Fridolfsson et al., 1998), and so all share ancestral ZW strata. The chromosome pair that was recruited to form the avian sex chromosomes is homologous to autosome 3 in *Alligator mississippiensis,* a member of Crocodilia, the closest extant clade to birds. *GHR* is just one gene among many that are homologous between avian Z and this crocodilian autosome. At the time of recruitment as sex chromosomes, Z and W would have had identical genetic content except for initial differences in the sex-determining region (Bachtrog, 2006; Behrens et al., 2024). For neoavians, *GHR* falls outside this region of ZW (see Zhou et al., 2014 for an overview of neoavian sex chromosome strata). Thus, ancestral birds would have had two copies of *GHR.* The sporadic appearance of *GHR-W* across the avian phylogeny (Figure 1), as well as the complete lack of any *GHR* copies outside of Z and W, is consistent with these genes being gametologs rather than paralogs.

### 4.2 Forces for Deletion or Retention of *GHR-W*

The fate of a gametolog is intertwined with the evolutionary trajectory of the sex chromosome on which it resides. Like the mammalian Y chromosome, the W chromosome has been subject to degradation over time, with cessation of recombination in increasingly large portions of the chromosome, enabling rampant insertion of repetitive elements, pseudogenization, and sequence loss (Zhou et al., 2014). For example, the chicken W chromosome is 14.15 Mb compared to the 86.74 Mb Z in a recent telomere-to-telomere assembly “GGswu1” (Z. Huang et al., 2023). Although some ZW homology has been maintained, allowing for recombination in PARs of varying size depending on the lineage, most of the W does not recombine in most avian clades (Zhou et al., 2014). W chromosome degradation is extensive in neoavians; for instance, outside the PAR, gene loss from W chromosome ranges from 88.12 - 97% in 15 neoavian and Galloanserine species examined in the literature (Sigeman et al., 2024; Zhou et al., 2014). The species retaining *GHR-W* do not have an unusually high rate of W sequence retention; for example, the Barn Owl (*Tyto alba*), which has *GHR-W,* has lost 93.32% of its W genes outside the PAR (Zhou et al., 2014). Further, in the immediate vicinity of the *GHR* gametologs, we find high levels of chromosomal degradation in high-quality neoavian genomes such as *Astur gentilis* (Figure 3B). Thus, if no forces favor retention, by far the most common fate for the long *GHR-W* gene (∼100 kilobase) should be loss. Retention of *GHR-W* in some neoavians thus suggests that selection may be acting to maintain these *GHR* copies on the W chromosome.

Additionally, as a gene that plays a critical role in development, *GHR* should be subject to strong purifying selection. Since this protein functions as a homodimer, having one faulty copy can effectively abolish the function of the working copy when they are dimerized (Derr et al., 2011). Therefore, retaining diverging copies of *GHR* outside the PAR is a risky proposition, with twice as many opportunities for mutations to create a dominant allele with a negative disease phenotype, again suggesting that positive selection may have played a role in maintaining *GHR* on W, despite the odds.

In contrast, the maintenance of *GHR-W* in certain paleognaths may owe more to this gene being found in the recombining PAR, which for example, spans much of the Emu (*Dromaius novaehollandiae*) Z and W chromosomes (Vicoso et al., 2013) including *GHR-Z* region. While less is known about the PAR in other paleognaths, which reportedly have greater levels of W degradation than *Dromaius* (Tsuda et al., 2007), in the Elegant Crested-Tinamou (*Eudromia elegans*) we find that *GHR-Z* and *GHR-W* are also located in a region with high sequence identity consistent with frequent recombination (see Figure 3A). Thus, it appears the maintenance of two *GHR* copies in birds is linked to the fate of the W chromosome as a whole, sometimes escaping widespread degradation (in the neoavians we examined), or persisting in a larger region of stability (in the paleognaths we examined).

### 4.3 *GHR-W* and Reverse Sexual Dimorphism

While we found *GHR-W* and RSD to be strongly correlated, *GHR*s are of course not the only genes to act in growth pathways. Many animals achieve larger female body size with an autosomal copy of *GHR*; e.g., the Eastern Painted Turtle, *Chrysemis picta* (Wilbur, 1975), which has *GHR* on chromosome 6 (GenBank: GCA_011386835.2). Likewise, we have not found evidence of *GHR-W* in chromosome-level genome assemblies in some birds with RSD, e.g. falcons. Our *GHR* copy number data is necessarily incomplete, as many taxa lack female genomes and, to date, no complete avian W chromosomes have been published (Zhao et al., 2025). Incomplete W chromosomes may be inadequate to determine the presence or absence of *GHR-W*. We look forward to further genome sequencing and assembly efforts in these taxa. In addition, further studies of taxa lacking sex-specific mass data will continue to reveal the contours of RSD in the avian tree of life. Even with the limitations of the available data, it is clear that *GHR-W* and RSD are highly correlated and co-occur in many avian taxa (Figure 4).

The majority of genomes in which we found evidence of *GHR-W* belong to accipitriform and strigiform birds of prey (hawks, owls and relatives). Both orders are well-known for larger female body size, i.e. RSD, and may well be sister groups phylogenetically. The correlation of *GHR-W* and RSD persists outside these taxa after correcting for oversampling of Accipitriformes (due to genome availability) and for phylogenetic structure. *GHR-W* and RSD appear together in multiple, deeply unrelated cladess in the avian phylogeny, for example, the paleognath orders Casuariiformes and Tinamiformes and sporadically throughout Neoaves. This repeated correlation is statistically unlikely to be the result of random chance (p < 0.000005), and could foreshadow a female-growth-specific role for a female-limited *GHR-W* gene.

While RSD is a rare phenomenon, it nevertheless has strong reproductive, evolutionary, and ecological implications for these species. Selective forces for larger females take a variety of forms across taxa with RSD, such as sexual selection in some Charadriiformes where females prefer smaller males due to their superior acrobatic mating flight displays (Blomqvist et al., 1997), selection on parental role specialization in Accipitriformes (Sonerud et al., 2014), and selection on resource partitioning in Accipitriformes (Bauld et al., 2022) (Slagsvold & A. Sonerud, 2007) (Krüger, 2005) and Suliformes (Zavalaga et al., 2007). The relevance of *GHR* to body size attainment in birds (Huang et al., 2016) may provide a modality for selection to act on the retention of two copies of this gene.

### 4.4 Potential Molecular Mechanisms of RSD

To affect female body size, *GHR-W* must be both intact and expressed. Our examination of assembled *GHR-W* and *GHR-Z* sequences in high-quality accipitriform genomes found an intact gene structure (exon-intron structure, splice sites, open reading frame). Additionally, RNAseq from female California Condors (*Gymnogyps californianus*) showed that both *GHR-W* and *GHR-Z* were expressed, albeit with *GHR-W* at a relatively lower level. As these transcripts were limited to adult fibroblasts, a tissue that generally has low *GHR* expression in vertebrates, it would be illuminating to explore data from more developmentally relevant tissues, such as liver, where *GHR* expression is key to body growth (Burnside et al., 1992).

*GHR-W* could affect female growth in two possible routes: via increased dosage or via female-specific changes. Neither Z chromosome is uniformly silenced in males, meaning that males can show higher expression of Z-linked genes (Davenport et al., 2023), and thus *GHR-W* could potentially bolster expression to match that of males. Having a female-limited *GHR* copy could also allow for sex-specific, copy-specific changes in gene regulation. This, or differences in the genomic environments of the two copies, could lead to combined *GHR-Z/GHR-W* expression in females exceeding male *GHR-Z* expression at key developmental stages where female growth accelerates vis a vis male growth.

Female-specific changes to the coding sequences that affect protein structure could impact *GHR* genes’ dual roles as a receptor, and also, if post-translationally modified, as a growth hormone binding protein that increases the longevity of growth hormone circulating in the blood (Schilbach & Bidlingmaier, 2015). Thus, evolution in the *GHR-W* gene sequence itself could have multiple avenues for increasing growth. Comparative data from species with RSD but without *GHR-W* could illuminate other sex-specific mechanisms for growth in birds.

## Supplemental Figure

**Supplemental Figure 1:**
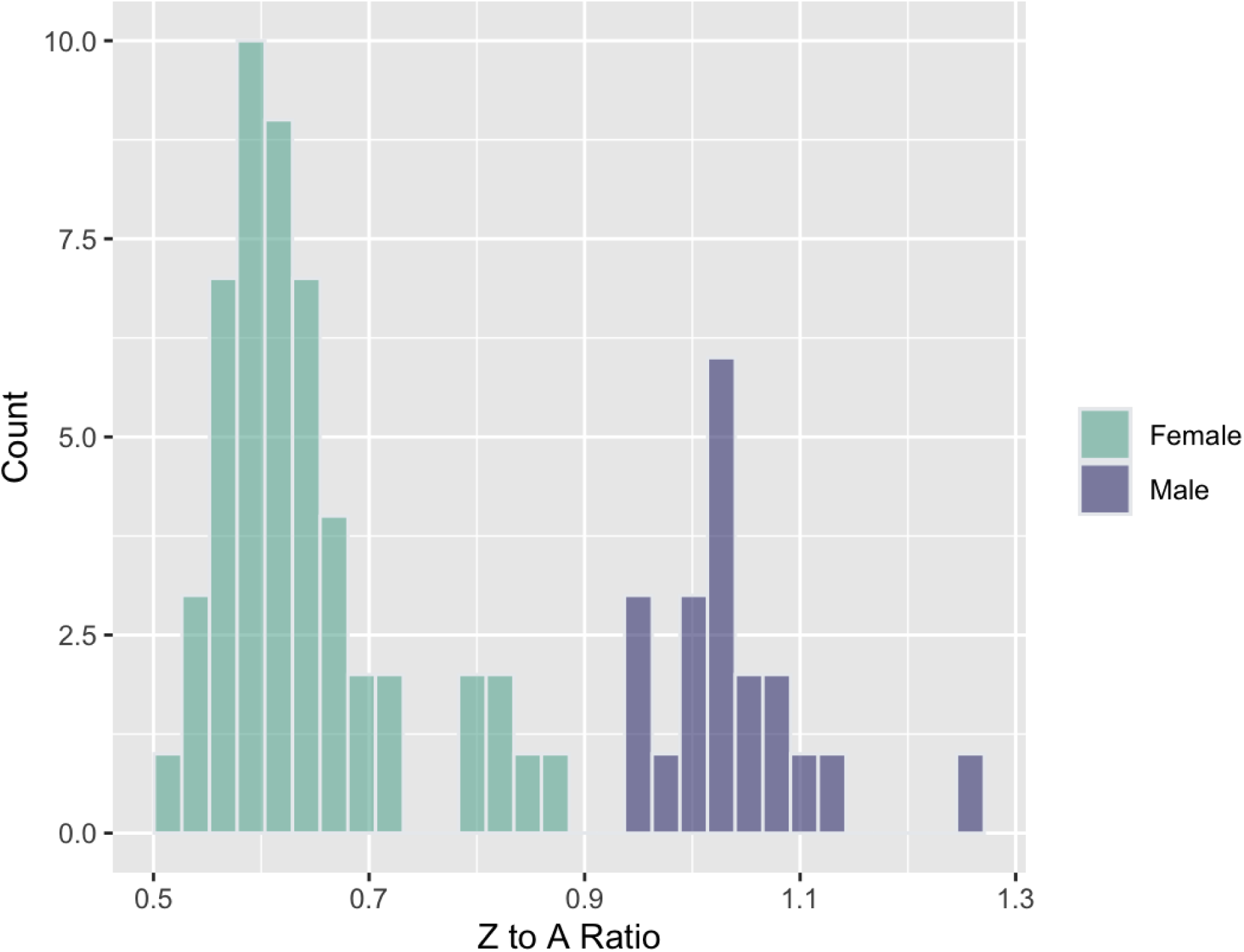
Histogram of Z:A chromosome coverage in birds that were sexed based on short read mapping (see Supplemental Table 2 for list of genomes). The ratio is calculated based on the mean coverage of the Z chromosome divided by the mean coverage of an autosome of similar length to the Z, as listed in Supplemental Table 2.

**Supplemental Figure 2:**
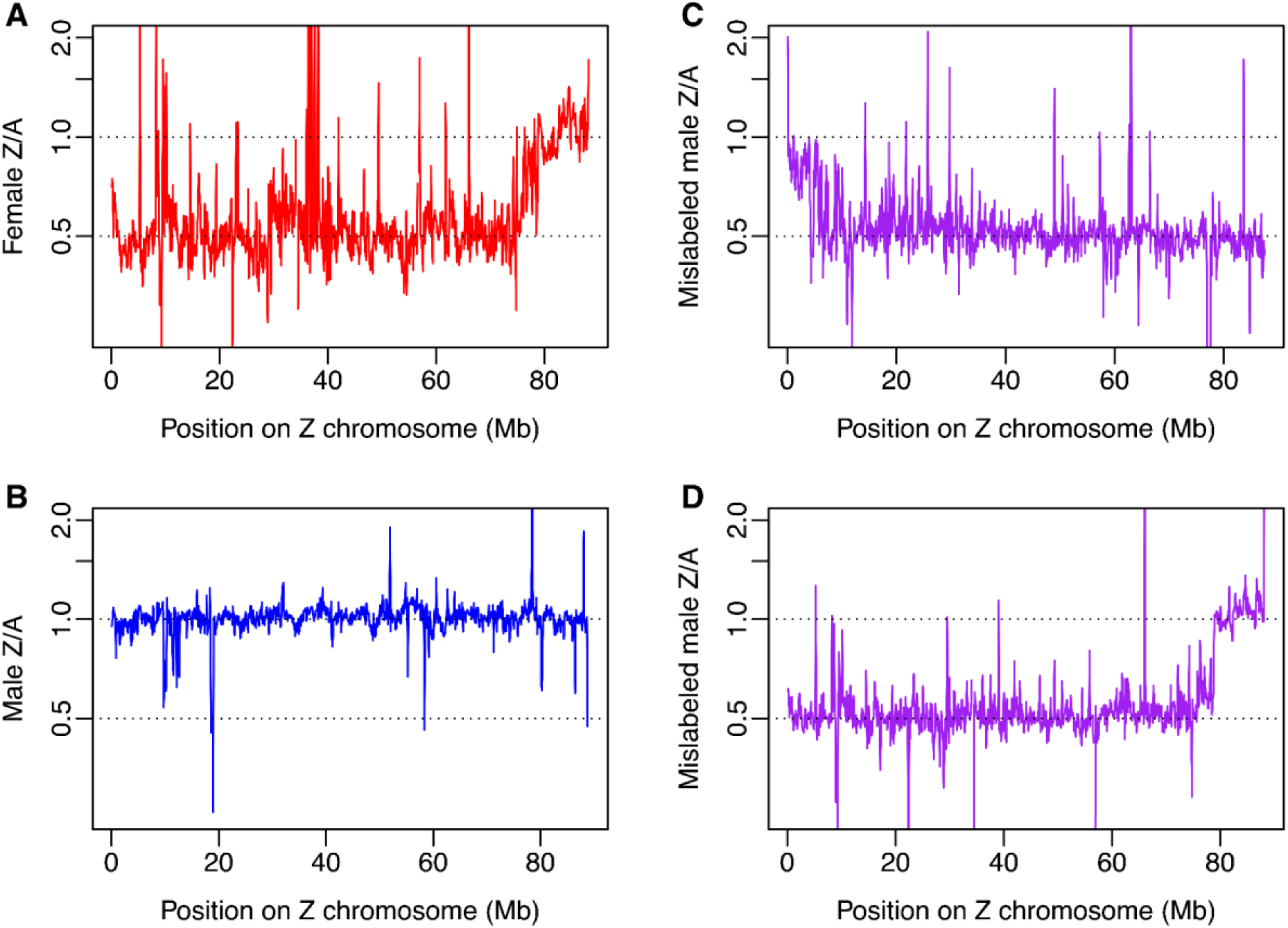
Ratio of Z to autosome (A) coverage in 100 kb sliding windows across Z chromosome. Z coverage is the mean over each window, and A coverage is the mean over the entire chromosome. The Z:A ratio displayed is calculated by dividing Z coverage by A coverage. ***2A.*** *Short sequence reads from a female* Accipiter striatus *genome* (GCA_027497855.1 - SRR17839736) *mapped to the chromosome-level* Aquila chrysaetos *genome (GCF_900496995.4). Coverage for the Z chromosome (GPC_000010101.1) divided by mean chromosome 1 (NC_044004.1) coverage is shown. As in many female genomes we examined, this mapping shows some elevated regions of Z coverage, driving the overall mean Z:A ratio to 0.78, well above the 0.5 ideal expectation for females. **2B.** Short sequence reads from a male* Aptenodytes forste*ri (GCA_000699145.1 - SRR1144986) mapped to the chromosome-level genome assembly for* Spheniscus humboldti *(GCA_027474245.1). Coverage for the Z chromosome (CM049923.1) divided by mean chromosome 5 (CM049891.1) coverage is shown. This mapping yields an overall mean Z:A ratio of 0.94, close to the ideal expectation of 1.0 for males. **2C & 2D.** Genomes that had hits for two copies of* GHR*and were labeled male in genome metadata or in the museum specimen listed in genome metadata, but we found had female Z:A ratios and hence relabeled as female in our data: **2C.*** Ibidorhyncha struthersii (GCA_013398815.1 - SRR9947056)*, mapped to* Pluvialis apricaria *GCA_017639485.1 with coverage for the Z chromosome (CM030044.1) divided by mean chromosome 4 (CM030010.1*) coverage is *shown; **2D.*** Milvus lineatus*, (GCA_035592195.1 - SRR25663121) mapped to* Aquila chrysaetos *genome (GCF_900496995.4). Coverage for the Z chromosome (GPC_000010101.1) divided by mean chromosome 1 (NC_044004.1) coverage is shown*.

## Author Contributions

All authors have contributed substantially to this work. Research conceptualization: S.A.R., M.J.B., P.L.F.J; Methodology: S.A.R., M.J.B., P.L.F.J; Data curation: S.A.R.; Data analysis: S.A.R., P.L.F.J.; Writing (original draft): S.A.R.; Writing (subsequent drafts, review, editing): S.A.R., M.J.B., P.L.F.J.

## Data Availability

All data referenced herein is available either directly as part of the supplementary materials or from NCBI RefSeq/GenBank via the genome/gene IDs listed in supplementary materials.

## Other

No ethics approval (e.g. animal or human experimentation), patient consent, permission to reproduce material from other sources, or clinical trial registration was needed for this work.

## Acknowledgements

We thank H.D. ‘Toby’ Bradshaw Jr. and Jennifer and Tom Coulson for helpful insights on raptor development.

